# Evolution of threespine stickleback dorsal spines via *hoxdb* gene regulation

**DOI:** 10.1101/2025.08.04.667941

**Authors:** Carlos Manuel Herrera-Castillo, Tanja Brechbühl, Antoine Fages, Andrew Donald Cameron MacColl, Laura L. Dean, Patrick Tschopp, Daniel Berner

**Affiliations:** Department of Environmental Sciences, Zoology & Evolution, University of Basel, Basel, Switzerland; School of Life Sciences, University of Nottingham, Nottingham, UK

## Abstract

The repeated emergence of similar phenotypes in independent populations is a widespread feature of evolution, yet the extent to which repeated evolution reflects shared molecular mechanisms, and the factors that determine this similarity, remain unclear. Here, we investigate the genetic basis of dorsal spine reduction in threespine stickleback fish, an adaptive trait that has evolved repeatedly in freshwater populations. Phenotypic analyses of crosses between a spine-reduced population from Scotland and the ancestral marine form reveal a major-effect locus, with the spine-reducing allele acting dominantly and influencing body patterning. Bulk segregant analysis maps this effect to a single region on chromosome VI that overlaps the *hoxdb* gene cluster. Transcriptomic analyses of larval stages show a gain of *hoxdb* expression in the developing spine region of spine-reduced fish, consistent with the dominance of the derived allele. Together, these results indicate dorsal spine reduction in threespine stickleback through cis-regulatory changes at *hoxdb*, a mechanism found to mediate dorsal spine evolution in a different species of stickleback fish too. These findings highlight the role of *hox* genes in adaptive divergence between natural populations within a species and provide a clear example of parallel evolution at the molecular level between species.

## Introduction

Understanding the molecular basis of adaptation remains a central challenge in biology, yet it is essential for addressing fundamental questions in evolutionary theory [1,2,3,4,5,6]. These include how phenotypic variation arises within populations, how novel variants spread through natural selection, as well as the impact and complexity of interactions between genomic loci in promoting adaptation, and the role of gene regulation in shaping adaptive phenotypes.

A particularly intriguing aspect of adaptation is repeated evolution – the independent emergence of similar phenotypes in response to comparable ecological conditions [1,7,8]. While such cases illustrate the potential predictability of natural selection at the phenotypic level, much remains to be uncovered about the molecular underpinnings of repeated adaptation. Elucidating these mechanisms is crucial for assessing what factors influence determinism in evolution [9,10,11,12,13,14]. For instance, specific features of genes and regulatory networks may increase the likelihood of their evolutionary reuse via mutational biases or inherent functional constraints [15,16,17,18]. In addition, demographic and population genetic factors such as divergence time, levels of gene flow and the availability of standing genetic variation, as well as the selective environment, may influence molecular repeatability [19,20,13,21,22,23,24,25,26,27].

Empirical data to evaluate these ideas remain limited but are increasingly generated by two complementary research strategies. The first approach involves ecological genomic studies aimed at identifying candidate regions under repeated selection by comparing genomic signatures across populations adapted to similar environments [26,28,29,30,31,32,33]. Relatively easy to implement, these methods have the additional advantage of potentially capturing the genetic basis of complex, polygenic adaptation. However, they generally face statistical challenges in reliably detecting true signals of selection and require follow-up studies to establish genotype-phenotype relationships [34,35]. The second approach focuses directly on phenotypes that have evolved repeatedly in independent populations, seeking to uncover their molecular basis at the gene or even mutation level [36,37,38,39,40,41,42,43,44]. Such phenotype-guided studies can provide deep insights into the repeatability of evolution but are typically labor-intensive, reductionist, and biased toward large-effect loci and easily measurable traits such as morphology [4,6,45]. Ecological genomic and phenotype-guided strategies thus offer complementary strengths and are both essential for a comprehensive understanding of the mechanisms driving repeated adaptation at the molecular level.

In this study, we investigate the molecular basis of repeated adaptation using a phenotype-guided approach in a well-established model system in evolutionary genetics, threespine stickleback fish (*Gasterosteus aculeatus*). Ancestrally marine, this species has repeatedly colonized freshwater habitats across its Holarctic range, giving rise to numerous cases of parallel evolution [46,47,48,49,50,51,52,21,53]. These encompass adaptive modifications to the external skeleton – including to the dorsal spines, after which the species has been named.

In stickleback and other spiny-rayed fishes, the dorsal fins are sub-divided into two separate evolutionary and developmental modules [54,55], with the unsegmented and heavily ossified spines of the anterior part serving for predator defense [56,57,58,59]. Most *G. aculeatus* populations possess three dorsal spines. The most posterior dorsal spine is almost universally present, and likely of limited ecological relevance. The two anterior and more prominent spines, however, show substantial variation between populations, with changes to both their size and number (Fig. 1a) [60,61,62,63]. Populations with reduced dorsal spines occur on the island of North Uist, Scotland [21,47,64,65,66]. In this system, reduction in the number of dorsal spines is considered an adaptive response to ion-depleted conditions, as these populations inhabit highly acidic and oligotrophic waters [62,64,65,66,67]. Additional evidence for local adaptation to limited resource availability includes a concurrent reduction in other skeletal elements such as lateral plates and the pelvic structure, as well as in overall body size [22,64,68].

**Fig. 1.**
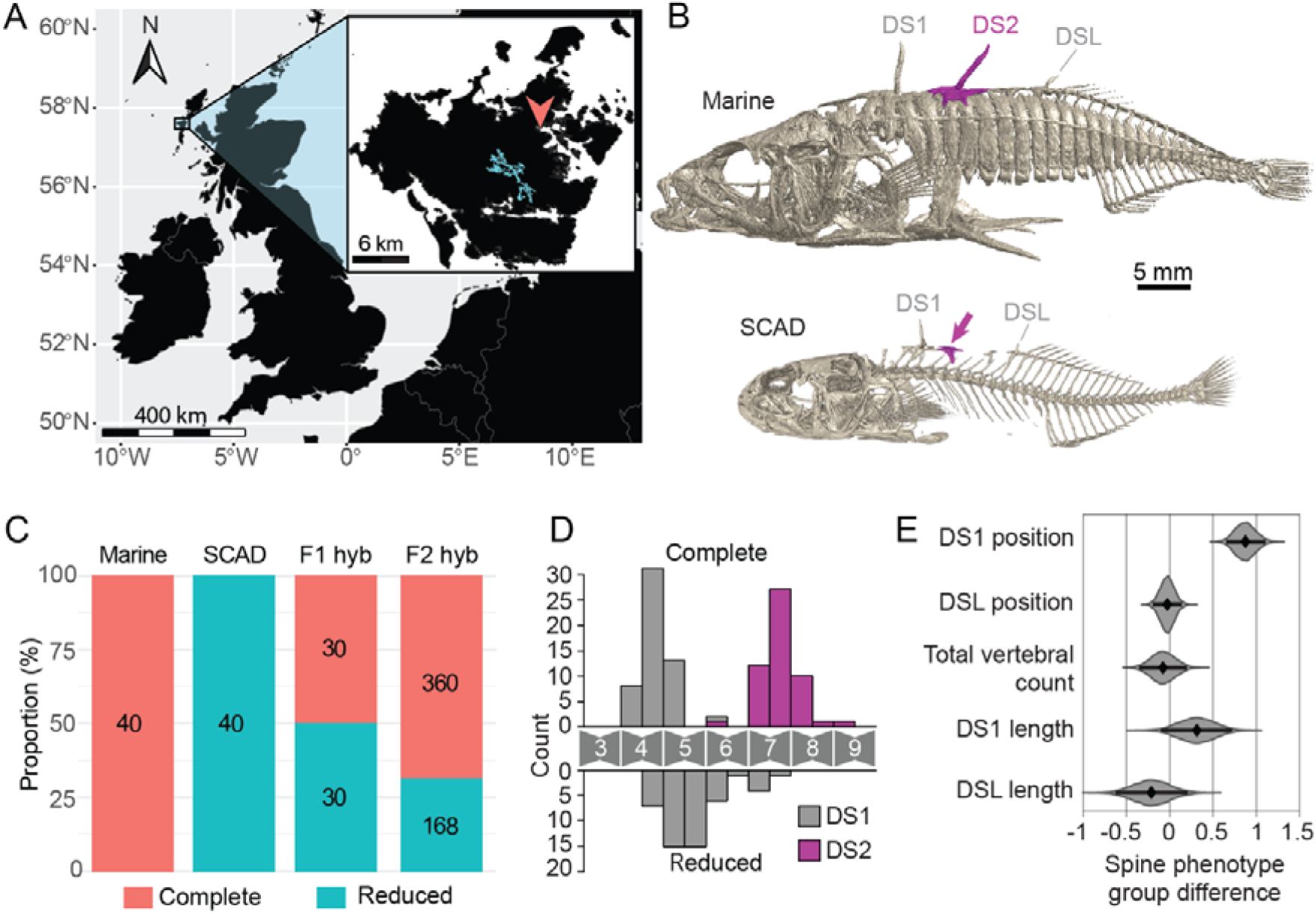
Origin and phenotypes of the study populations and their crosses. **A)** Geographic situation of North Uist island (close up) in the Outer Hebrides, Scotland. Lake Scadavay (SCAD) is shown in blue and the sampling site of the marine ecotype is indicated by the red arrowhead. **B)** X-ray image of a representative individual from each ecotype, visualizing only bony structures. The spine-reduced SCAD ecotype typically lacks the second dorsal spine (DS2) consistently present in the completely spined marine ecotype, but often exhibits the skeletal base (pterygiophore) that normally supports that spine (arrow). **C)** Proportion of completely spined and spine-reduced individuals in the pure parental experimental lines, and in the F1 and F2 hybrid crosses derived from these parents. **D)** Position of the two major dorsal spines in completely spined individuals, and of the first dorsal spine in spine-reduced individuals from the F2 hybrid cross, using the vertebrae as a reference scale. **E)** Difference between completely spined and spine-reduced F2 hybrid individuals in the position of the first (DS1) and last (DSL) dorsal spine, in total vertebral count, and in the body size-adjusted length of DS1 and DSL. For the upper three traits, the scale refers to vertebrae, whereas the spine lengths represent standardized data. For all traits, the diamond indicates the point estimate, the thick black line the associated 95% bootstrap compatibility interval, and the violins the full bootstrap distribution. The difference is always expressed as SCAD minus marine.

Despite these striking phenotypic changes, the molecular basis of dorsal spine reduction in threespine stickleback remains unknown. In this study, we address this gap by combining developmental, genomic, and transcriptomic analyses in two populations from North Uist, an ancestral marine and a spine-reduced lacustrine population. This reveals that spine reduction is driven by a developmental patterning gene cluster also found to mediate dorsal spine evolution in a different stickleback species, thus providing a striking case of repeated evolution.

## Results and Discussion

### Phenotypic analysis suggests a large-effect locus with a dominant derived allele influencing dorsal spine patterning

To investigate the genetic basis of dorsal spine reduction, we focused on a lacustrine population of threespine stickleback from Loch Scadavay (hereafter “SCAD”) on North Uist (Fig. 1A) [64,69]. This freshwater ecotype is generally spine-reduced (Fig. 1B bottom), with approximately 90-95% of the individuals lacking the second dorsal spine. As a representative of the ancestral ecotype from which the SCAD population originates, we considered marine stickleback (Fig. 1B top) sampled from a coastal estuarine breeding site on the same island (hereafter “marine”, Fig. 1A). From field-caught individuals, we established pure laboratory lines for both ecotypes, as well as multiple F1 and F2 intercrosses (hereafter “hybrids”). In addition, a single-parent F4 hybrid cohort was reared in a natural pond environment (“pond cohort”). To assess the inheritance of dorsal spine reduction, individuals from all lines were scored for the presence or absence of the second major dorsal spine, classifying them as completely spined or spine-reduced.

All laboratory-reared marine individuals were completely spined, whereas all SCAD individuals were spine-reduced (Fig. 1C). Under shared controlled conditions, the pure lines thus faithfully recapitulated the dorsal spine differences observed between wild SCAD and marine stickleback [21,22,64,69,70]. This supports a strong genetic basis for this ecotype divergence, rather than an immediate environmental effect on development. The F1 hybrids displayed approximately equal proportions of completely spined and spine-reduced phenotypes, whereas the F2 hybrids showed roughly one-third spine-reduced individuals (Fig. 1C). Segregation analysis and simulations indicated that these phenotype distributions are consistent with a single locus of large effect, with dorsal spine reduction caused by a dominant derived allele segregating at an estimated frequency of 60-70% within the SCAD population (S1 Analysis).

Spine development and diversification in acanthomorph fishes is linked to anterior-posterior dorsal fin patterning [54,55,71], including the distribution of the skeletal fin bases (pterygiophores) along the underlying vertebral column [72,73]. Accordingly, we first examined whether spine reduction in our F2 hybrid population coincided with changes in dorsal spine position relative to vertebral position. We found that spine-reduced fish exhibited a posterior shift of the first dorsal spine (DS1) by nearly one vertebra, and that the position of this spine was more variable than in completely spined individuals (Fig. 1D, E). By contrast, we observed no relevant difference between the two phenotypic classes in the position of the small last dorsal spine (DSL) marking the onset of the dorsal fin, nor in total vertebral number (Fig. 1E). Likewise, we found no substantial or consistent difference in the length of DS1 and DSL; small possible effects in either direction were fully compatible with our data (Fig. 1E). Together, these results are consistent with variation in dorsal spine number being mediated by a developmental patterning locus specifically affecting anterior spine positioning.

### Divergence in dorsal spine formation characterized by developmental series

To characterize the ontogenetic origin of dorsal spine number variation, we generated postembryonic developmental series for SCAD and marine laboratory lines. Larvae were sampled over the first 30 days post hatching (dph) and subjected to cartilage (Alcian Blue) and bone (Alizarin Red) staining.

In the marine ecotype, development of DS2 was initiated by the formation of a cartilaginous pterygiophore scaffold, first detectable as a small focal condensation at approximately 9 dph (Fig. 2A). This cartilage element subsequently extended toward the vertebral column and then ossified to give rise to the final pterygiophore (Fig. 2B). The spine itself emerged atop the still cartilaginous pterygiophore and ossified directly, without an apparent cartilaginous precursor (Fig. 2B). By 22 dph, all marine individuals possessed fully ossified pterygiophores and three dorsal spines (Fig. 2A, B). This developmental sequence is consistent with that described for other completely spined populations of threespine stickleback [74,75].

**Fig. 2.**
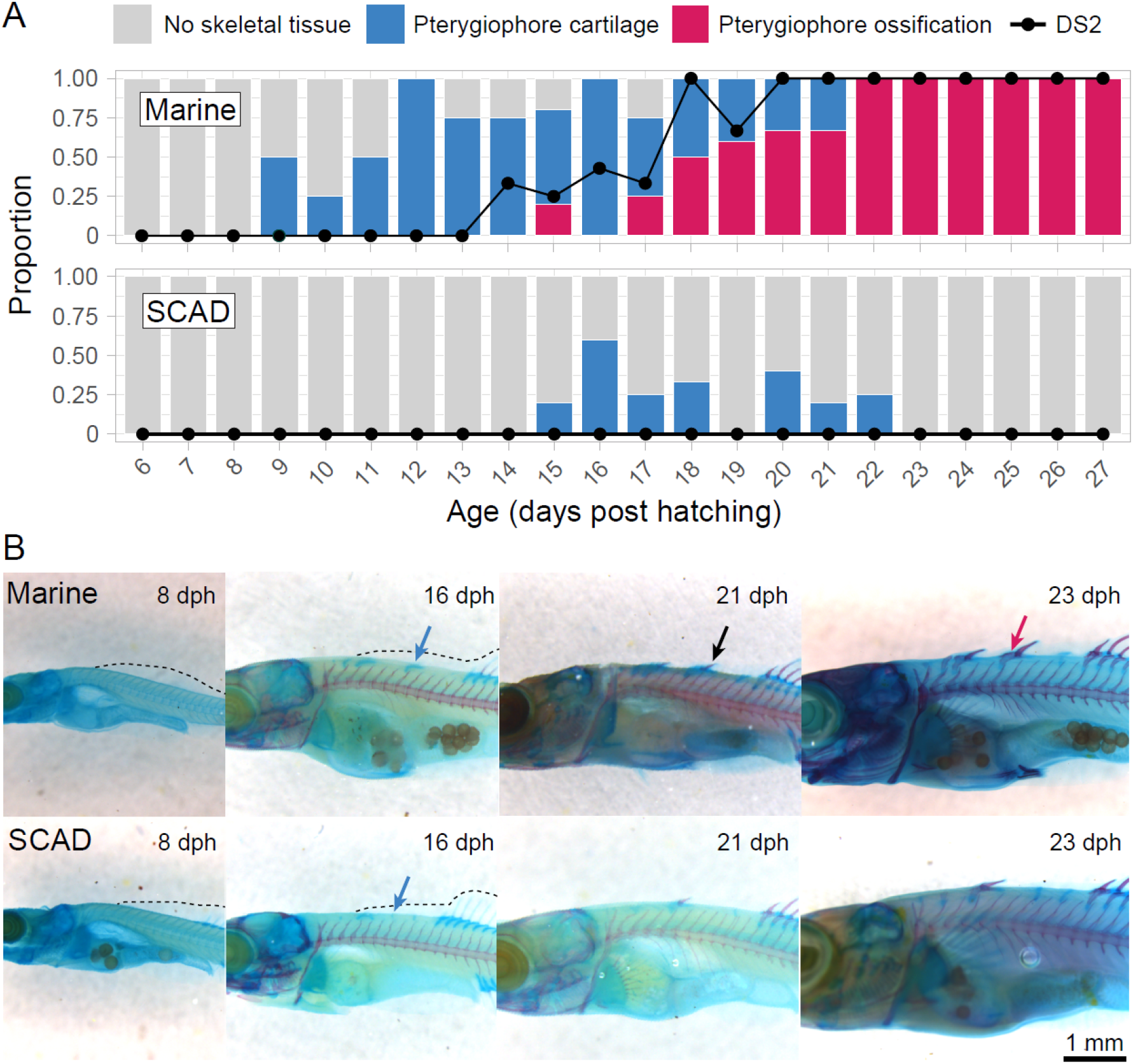
Dorsal spine development, or the lack thereof, in marine and SCAD threespine stickleback. **A)** Developmental series of larval cartilage and bone formation, focusing exclusively on elements of DS2. The vertical bars indicate the proportion of individuals in specific stages of the development of the skeletal base (pterygiophore) on top of which DS2 may later form. The development of DS2, which is not preceded by cartilage, is illustrated by the black line. Note that the proportion of individuals with spine ossification does not increase strictly monotonically because the data come from different individuals slightly varying in developmental stage. **B)** Representative cartilage and bone double-stained marine and SCAD larval individuals in key stages of pterygiophore and DS2 development. Blue and red arrows indicate cartilage and bone elements of DS2’s pterygiophore, while the black arrow indicates the ossification of DS2 itself. For the marine population, the panels illustrate, from left to right, the stage prior to any development of the focal structures, pterygiophore cartilage formation, incipient spine ossification, and pterygiophore ossification. In SCAD individuals, DS2 development does not progress beyond weak pterygiophore cartilage formation, while DS1 develops normally. The dashed line indicates the median fin fold (mostly regressed outside the fin domain in the later stages).

Dorsal spine development in SCAD larvae differed from the marine trajectory in important ways (Fig. 2A, B). First, the initiation of the DS2 cartilaginous pterygiophore was either delayed by up to six days or failed to occur altogether. This structure also did not extend or ossify into a mature pterygiophore in any of our focal experimental fish. However, haphazardly inspecting adult SCAD specimens sometimes revealed an ossified pterygiophore posterior to the one supporting DS1 (e.g., Fig. 1B, arrow), suggesting that some pterygiophore development beyond 22 dph may have been overlooked. Finally, and most importantly, SCAD individuals showed no evidence of DS2 formation at any developmental stage examined.

### Genome scans suggest the hoxdb gene cluster as a candidate locus

To identify the large-effect locus suggested by our phenotypic analyses (Fig. 1C and Analysis S1), we performed a bulk segregant analysis on 469 F4 hybrid individuals from the pond cohort. Fish were phenotyped for dorsal spines and assigned either to a completely spined group (DS2 present) or a spine-reduced group (DS2 lacking). Using pooled whole-genome sequencing, we scanned all chromosomes for regions showing pronounced allele frequency differentiation between the two groups. This analysis revealed a single region at the right end of chromosome VI clearly associated with dorsal spine phenotype (Fig. 3A,B). Several additional narrow peaks driven by only a few SNPs were also detected (Fig. 3A), which we interpret as technical artifacts; given the limited opportunity for recombination in an F4 cohort, any true genotype-phenotype association is expected to span megabases comprising tens of thousands of SNPs. As shown by the observed peak on chromosome VI (Fig.3B), this also holds for the high-recombination chromosome peripheries characteristic of threespine stickleback [76].

**Fig. 3.**
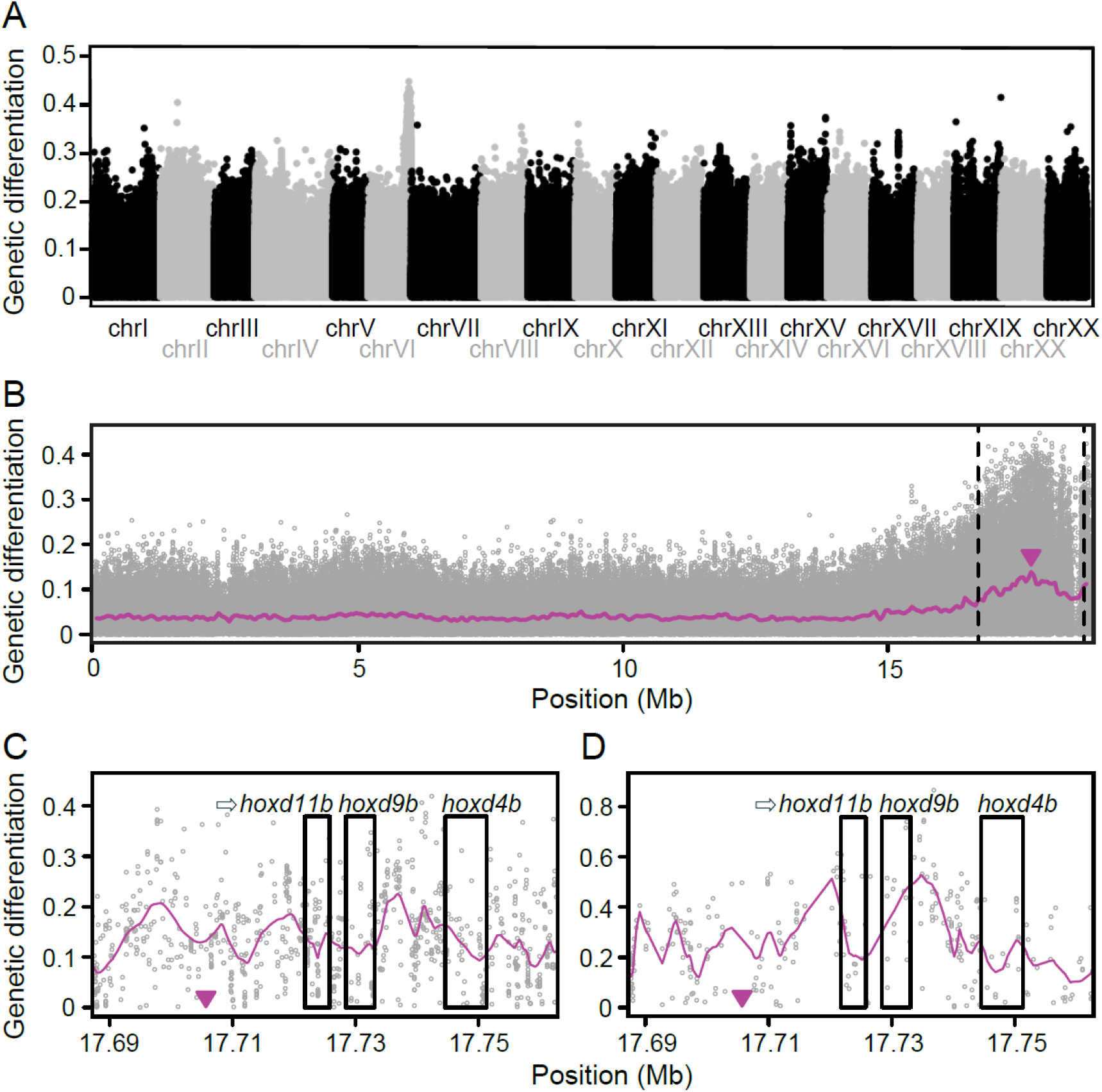
Genome scans for the genetic basis of divergent dorsal spine development. **A)** Genome-wide genetic differentiation across the 21 threespine stickleback chromosomes revealed by bulk segregant analysis comparing completely spined and spine-reduced individuals from the F4 hybrid pond cohort. Dots indicate differentiation at individual SNPs, expressed as the absolute allele frequency difference (AFD). **B)** Same analysis as in A), but restricted to chromosome VI, which harbors the only well-supported differentiation peak. The purple line represents SNP differentiation, smoothed using 100-kb sliding windows with 50-kb overlap. The purple arrowhead marks the midpoint position of the sliding window with the highest median AFD genome-wide. Dashed vertical lines delimit the 2-Mb region used as the search interval for genes showing differential expression between marine and SCAD fish in the bulk RNA sequencing analysis shown in Fig. 4. **C)** Fine-scale view of bulk segregant differentiation focusing on a 75-kb region on chromosome VI encompassing both the peak position identified in B) (purple arrowhead) and the *hoxdb* gene cluster. The arrow before the *hoxd11b* label indicates the transcriptional orientation of the entire gene cluster. **D)** AFD across the same region as in C), observed in the comparison of SCAD and completely spined North Uist freshwater stickleback based on pooled sequencing. In C) and D), the purple lines indicate LOWESS-smoothed AFD.

Inspecting the gene annotation within the chromosome VI peak identified a compelling candidate locus: the *hoxdb* cluster, containing genes *hoxd11b, hoxd9b, hoxd4b*, all of which are located within 50 kb of the estimated differentiation peak (Fig. 3C). *Hox* genes encode highly conserved developmental regulators that define regional identities and play central roles in axial and appendicular patterning across vertebrates and bilaterians at large, including for the formation of median fins in teleost fish [55,77,78]. These transcription factors are usually found tightly organized in genomic clusters, with the relative positioning inside these loci having important implications for their spatial and temporal gene expression [79,80].

To obtain independent population-level support for the *hoxdb* region as a candidate locus, we conducted a population differentiation scan using pooled whole-genome sequencing data from natural stickleback populations. This analysis compared a single DNA pool of 130 spine-reduced SCAD individuals with a pool of 100 individuals sampled from five freshwater populations from North Uist harboring exclusively completely spined phenotypes. Assuming that the latter populations are fixed for the ancestral recessive allele, we expected the genomic region harboring the *hoxdb* cluster to exhibit allele frequency differences of around 0.6-0.8 between the two groups (S1 Analysis). Because this analysis leveraged haplotype diversity in natural populations rather than a controlled cross, it provided substantially higher physical resolution than the bulk segregant analysis. Allele frequency differentiation consistent with our expectation was indeed detected in genomic segments immediately upstream of *hoxd11b* and downstream of *hoxd9b* (Fig. 3D). While this population-level scan does not by itself establish a direct association between allele frequency differentiation and dorsal spine phenotype, observing the predicted differentiation signal within the chromosome VI region identified in the bulk segregant analysis nonetheless supports a role for the *hoxdb* locus in dorsal spine evolution across natural stickleback populations.

### Gene expression analysis indicates regulatory evolution at hoxdb

*Hox* genes play central roles in the embryonic patterning and differentiation of multiple vertebrate organ systems [81,82]. This makes changes in their coding sequences often highly pleiotropic and potentially detrimental. We therefore hypothesized that dorsal spine evolution in our stickleback system is driven by *cis*-regulatory variation at the *hoxdb* cluster, that is, by non-coding polymorphisms modulating the timing and/or spatial pattern of gene expression. We explored this prediction using a transcriptomic time series of tissue samples that were micro-dissected from the developing dorsal spine region of multiple pure marine and SCAD individuals. The samples were collected at four larval ages (10, 13, 16, and 19 dph), thus spanning the time period and anatomical location of major developmental divergence between the two ecotypes (Fig. 2).

Principal component analysis (PCA) based on the expression of the top 5000 variables genes in the genome-wide transcriptomic data revealed a clear separation between the two ecotypes, but not by larval age (S1 Figure A). Combined across the age classes, differential expression between marine and SCAD individuals was found in 623 of these genes (S1 Figure B), and gene ontology enrichment analysis revealed that they were enriched for terms related to extracellular structure organization and remodeling (S1 Figure C), reflecting the observed phenotypic differences in dorsal spine development. Beyond these genome-wide expression signals, focus on the candidate region showed that among all genes located within a 2 Mb window centered on the differentiation peak identified in the bulk segregant analysis (Fig. 3B), substantial expression differences were restricted exclusively to the *hoxdb* cluster, with the strongest effects observed for *hoxd11b* and *hoxd9b* (Fig. 4A).

**Fig. 4.**
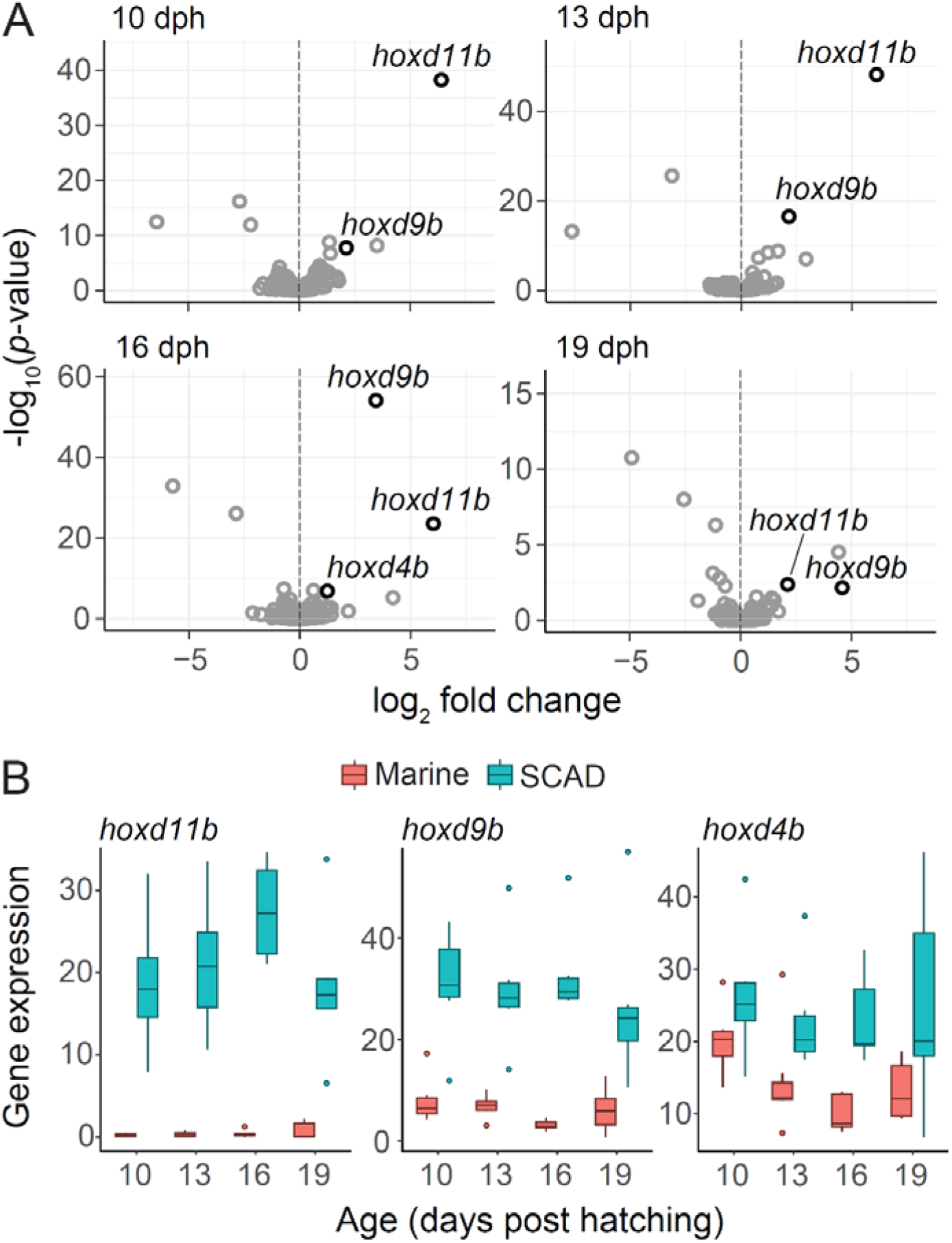
Differential expression of *hoxdb* genes between marine and SCAD stickleback larvae. **A)** Differential expression between ecotypes, considering all genes located on chromosome VI. The x-axis represents the log_2_ fold change in expression between the two ecotypes, with positive values being reflecting higher expression in SCAD population relative to the marine one. Data are shown separately for each of the four focal age classes (10, 13, 16 and 19 days post hatching), with each gene represented by a single point. Genes located within the 2-Mb search interval shown in Fig. 3B and qualifying as strongly differentially expressed (log_2_ fold change ≥ 1, −log_10_(*p*-value) ≥ 2) are highlighted in black; all of these represent *hoxdb* genes. **B)** Boxplots showing normalized (CPM) expression levels of the three focal *hoxdb* genes in marine and SCAD ecotypes across the four developmental time points. All analyses are based on 30 total tissue samples per ecotype and age class combination, grouped into six biological replicates.

The concordance between strong sequence differentiation (Fig. 3) and differential expression of *hoxdb* genes between spine phenotypes provides compelling evidence that dorsal spine evolution is related to regulatory variation at this locus [3,5]. Consistent with this view, differentiation in the population-level comparison proved most pronounced in the noncoding region between *hoxd9b* and *hoxd4b* (Fig. 3D). Interestingly, the *hoxdb* genes were overexpressed in spine-reduced fish relative to the marine ecotype (Fig. 4B), suggesting that the underlying regulatory element may normally function as a silencer and that the derived SCAD variant disrupts this repression, resulting in elevated *hoxdb* transcription. Indeed, in mice, the central intergenic region of the *HoxD* cluster has been reported to act as a regulatory buffer, by preventing anterior enhancers from ectopically activating posterior genes [83,84]. Loss of repression would also provide a plausible explanation for the dominance of the variant segregating within SCAD: owing to nonlinear genotype-phenotype relationships, a single copy of a gain-of-expression allele is often sufficient to produce a phenotypic effect, such that dorsal spine reduction would be expected even in heterozygous individuals [85,86,87].

### Hoxdb is involved in repeated dorsal spine evolution in stickleback fish

Dominance at *hoxdb* contrasts with the phenotypic effects of large-effect derived alleles underlying ecologically important trait variation identified so far in threespine stickleback, including the derived alleles at *eda* and *pitx1* associated with lateral plate and pelvic reduction [88,89,90]. This distinction has two important implications. First, because the derived *hoxdb* spine reduction allele is expressed in heterozygotes, it is immediately visible to selection even when present in a single copy, potentially enabling rapid adaptive responses under favorable ecological conditions. Conversely, the absence of masking in heterozygotes should lead to efficient purging when the phenotype is maladaptive. Reduced dorsal spines are likely strongly disadvantageous in the marine environment, where sticklebacks are typically heavily armored; to our knowledge, marine individuals lacking the second dorsal spine have not been reported. Unlike recessive freshwater-adaptive alleles, which can persist cryptically in marine standing genetic variation and thereby facilitate repeated adaptation across populations, the derived *hoxdb* allele is therefore unlikely to be maintained in the ancestral marine gene pool.

These properties make *hoxdb* a strong candidate for repeated evolution of dorsal spine number but suggest that geographically distant populations may evolve this phenotype through independent mutations rather than shared standing variation. Supporting this idea, dorsal spine number evolution involving the *hoxdb* cluster has also been documented in the fourspine stickleback, *Apeltes quadracus* [71,91], thus representing a striking case of repeated evolution across species separated by tens of millions of years. However, repeated evolution via *hoxdb* may also occur within threespine stickleback; in Boulton Lake, British Columbia, Canada – an acidic habitat similar to SCAD [61] – most individuals likewise lack the second dorsal spine [60,63]. Although the exact genetic basis of spine reduction in this population remains unclear, cross data [71] indicate segregation of a derived allele with a similarly large and dominant phenotypic effect (S1 Analysis).

Additional threespine stickleback populations with dorsal spine reduction occur in close geographic proximity to SCAD. Comparative genetic analyses of spine-reduced populations across different geographic distances could therefore provide a valuable opportunity to examine to what extent the probability of repeated evolution via independent mutation versus alleles identical by descent is influenced by the phenotypic effect of the underlying locus.

### Conclusions

Our study reveals adaptive skeletal evolution in a European stickleback population via variation in a *hox* gene cluster. Spine evolution in our stickleback system is thus not based on a locus involved in localized dorsal spine skeletogenesis, but rather on alterations in body patterning. *Hox* genes are well known to determine patterning in the embryonic stage, and to be responsible for major developmental evolution at relatively high taxonomic level. The finding of a *hox* locus influencing larval spine development and driving ecologically relevant phenotypic variation within and among stickleback populations highlights the diverse transcriptional regulation and evolutionary roles of *hox* genes.

The repeated use of the *hoxdb* cluster across stickleback species suggests that spine evolution is biased toward this locus. We speculate that one element of this bias is the dominant phenotypic effect of *hoxdb* variants, making them immediately visible to natural selection. Uncovering the specific mutations underlying dorsal spine reduction in multiple populations of threespine stickleback will allow exploring the eco-geographic distribution of the underlying variants, facilitate experiments quantifying the fitness consequences of spine variation across different habitats, and promote our understanding of adaptive evolution in gene regulation.

## Materals and Methods

### Study system and experimental lines

Threespine stickleback were sampled from two populations on North Uist Island, Scotland: a lacustrine freshwater ecotype from Loch Scadavay (hereafter “SCAD”; 57°35′44.0″ N, 7°15′13.0″ W) and a fully spined marine ecotype from a coastal lagoon breeding site at Loch an Duin (57°38′32.96″ N, 7°12′31.76″ W). Our SCAD sample considered spine-reduced individuals only, although completely spined fish occur in that population at low frequency (S1 Analysis). Laboratory lines for both ecotypes were initiated in spring 2021 using embryos produced by *in vitro* crosses of the field-caught adults, with 10–12 independent crosses per population, each using unique parental individuals. Clutches were dissociated into individual eggs and shipped to the University of Basel at 5°C in 50-mL tubes containing a dilute methylene-blue solution to prevent infection. Upon arrival, eggs were transferred to 15-L tanks maintained at 16 °C under a long-day photoperiod (16 h light : 8 h dark), which is our standard rearing regime. Following established rearing and breeding protocols [92], this first laboratory cohort was raised to adulthood and used to generate a second laboratory cohort comprising pure lines (36 unique families per ecotype) and between-ecotype F1 “hybrids” (five unique families). Adults from these lines were subsequently used to produce a third laboratory cohort, including pure lines and F2 hybrids (40 families), the latter generated by haphazard intercrossing among individuals from the F1 hybrid families.

In addition to these laboratory-derived lines, our study includes F4 hybrids originating from a single cross performed in spring 2015 between a SCAD male and a marine (Loch an Duin) female. This hybrid population (“pond cohort”) was established in 2016 by releasing 101 adult F1 hybrids into a pond (previously without stickleback) near the University of Nottingham, England, and reproduced naturally for at least two generations; sampling occurred in 2018.

### Phenotypic characterization of the laboratory lines

To gain insights into the inheritance of the loss versus presence of the dorsal spine in our study system, we sampled a total of 40 adult individuals per ecotype from the first and second laboratory cohort, plus 60 F1 hybrids and 528 F2 hybrids from the second and third cohort. These individuals were inspected under a stereomicroscope and classified as completely spined (i.e., DS1-DSL present) or spine-reduced (DS2 missing, just DS1 and DSL present) (Fig. 1B). For visualization of the pure ecotypes, a representative individual from SCAD and from the marine ecotype were fully dried and subjected to a 180° X-ray micro-computed tomography (μCT) scan on a Bruker Skyscan instrument with a 1-mm aluminum filter, 3300 ms exposure time, 50 kV voltage, 800 μA current, and 0.8° rotation steps.

Given the marine population is likely fixed for the ancestral allele, the high proportions of spine-reduced individuals in both our F1 and F2 hybrid cross clearly indicated a major effect locus with dominant phenotypic effect within SCAD. To explore the frequency *p* of that derived, dominant allele within SCAD, we first ran individual-based simulations mimicking our laboratory breeding scheme to produce hypothetical proportions of ancestral (completely spined) individuals in our F1 and F2 hybrid crosses as a function of different values of *p*. We here aimed to approximate our actual experimental conditions by modeling: full dominance for the derived allele; 20 pure parental individuals initiating the laboratory lines, with the marine ecotype fixed for the ancestral allele and the SCAD population sampled stochastically (i.e., with replacement) from Hardy-Weinberg genotype proportions defined by *p*; 60 total F1 hybrid individuals resulting from five F1 families with family sizes of 6, 11, 13, 13 and 17 individuals produced by random pairing of unique marine and SCAD parents; 530 F2 hybrid individuals obtained by random pairing of F1 parents. For each of the 10,000 simulation runs thus performed, *p* was drawn at random between 0 and 1 with uniform probability, and the proportion of the ancestral phenotype within the F1 and F2 cohort was recorded. The most plausible frequency of the derived allele in SCAD given our simulated phenotype proportions was determined by Approximate Bayesian Computation (ABC; [93]), using the observed proportions of the ancestral phenotype in the F1 and F2 hybrid cohort (i.e., 0.5 and 0.68, Fig. 1C) as estimation targets. We performed 500 ABC estimation runs with the neural network algorithm and a tolerance of 0.05 (tolerances of 0.01, 0.02, and 0.1 produced very similar results), and evaluated the distribution of the estimated medians across the replicate runs.

Our second approach to estimate *p* used the approximate proportion of the ancestral phenotype in the natural SCAD population, as estimated from multiple rounds of field sampling. This proportion is approximately 5-10% (A. MacColl, personal estimation). Assuming full dominance of the derived allele and genotypes in Hardy-Weinberg equilibrium, only individuals homozygous for the ancestral allele express the ancestral phenotype; thus, the observed phenotypic frequency corresponds to the homozygous ancestral genotype frequency, (1 - *p*)^2^.

To assess whether the genetic factor underlying dorsal spine reduction also affects other patterning and spine traits, we analyzed individuals from the F2 hybrid cohort. Flatbed X-ray images (taken with Kubtec MOZART Supra Specimen Tomosynthesis System) were obtained for 49 spine-reduced and 53 fully spined individuals. From these images, we recorded the presence and positional location of all dorsal spines. Spine position was determined by projecting the spine articulation points perpendicularly onto the vertebral column, with positions scored at a resolution of 0.5 vertebra (i.e., anterior or posterior half of a given vertebra), where vertebral position 1 corresponded to the first fully visible vertebra. The vertebral column was used solely as a positional reference scale, without assuming any fixed correspondence between spine formation and body segmentation. For each individual, we also counted the total number of vertebrae, starting with the first fully visible vertebra and excluding the enlarged terminal vertebral element associated with the hypural plate. Differences between spine-reduced and fully spined groups were quantified using compatibility intervals, defined as the central 95% percentiles of bootstrap distributions generated by resampling each group 10,000 times and recalculating the mean group difference in each iteration [94].

The F2 hybrid cohort was also used to test for a positive association between dorsal spine number and dorsal spine length, as previously reported for another spine-reduced threespine stickleback population [71]. Macro-images with an embedded reference scale were obtained for 37 spine-reduced and 39 completely spined individuals using a CMEX-5f ocular camera (Euromex) mounted on a stereomicroscope. DS1 and DSL lengths were measured from these images using ImageJ [95].

To facilitate comparison, measurements for each spine were mean-centered across all individuals and standardized. Because dorsal spine length scales with overall body size, phenotype group differences for each spine were estimated using linear models with phenotype group as the main predictor and body size as a covariate [96], after confirming that the phenotype x body size interaction was negligible. Body size was quantified as scores along the first principal component (PC1; 92% of variance explained) from a principal component analysis of standard length and cube-root transformed body mass of the preserved specimens. Compatibility intervals for spine length differences between the phenotype groups were obtained by bootstrapping as described above.

### Developmental series

A median of four larval individuals per ecotype were sampled from the first laboratory cohort daily from 6 to 35 days post hatching (dph), covering the Swarup stages 28-31 [74]. All larvae were then subjected to simultaneous whole-body staining with Alcian blue for cartilage and Alizarin red for bone, following the acid-free double staining protocol described in Walker & Kimmel [97]. The stains were then photographed using a Leica stereomicroscope (Leica M205 FA) with a mounted digital camera (Leica DFC310 FX) and the Leica Application Suite, v.3.7.0 (build 681) to characterize differences between the ecotypes in the presence or absence of cartilage and bone elements, as well as the timing of their appearance.

### Bulk segregant analysis

To identify genomic regions associated with dorsal spine reduction, we used the pond F4 hybrid cohort to scan the stickleback genome for loci showing strong differentiation between spine phenotypes, following the pooled-sequencing strategy of Laurentino et al. [98]. A total of 469 ethanol-preserved adults were stained with Alizarin red (protocol provided in S1 Methods) to visualize skeletal elements and scored for dorsal spine development. This screening identified 163 spine-reduced individuals, which were assigned to one pool, and 306 completely spined individuals, which formed the alternative pool. Genomic DNA was extracted from each individual using the Quick-DNA Miniprep Plus Kit (Zymo Research) following the manufacturer’s protocol, with minor modifications as described in Laurentino et al. [98]. DNA concentrations were quantified using a Qubit fluorometer with the Broad Range kit (Invitrogen, Thermo Fisher Scientific, Wilmington, DE, USA). DNA from individuals within each phenotype group was then pooled in equimolar proportions, and the two pooled libraries were sequenced to 100 bp paired-end reads on an Illumina NovaSeq S4 flow cell at the Genomics Facility Basel (D-BSSE, ETH Zürich). Median genome-wide read depth was 170 for the spine-reduced pool and 199 for the fully spined pool.

The raw sequences were aligned to the fifth-generation assembly of the threespine stickleback genome (447 Mb; [99]). The locus harboring the *pitx1* gene (378 kb in length), available from GeneBank (accession: GU130435.1) but missing from the reference genome, was included as a separate chromosome. The sequences were aligned by using Novoalign (v.3.02.00; http://www.novocraft.com; parameter settings given in Supplemental Code). Using the Rsamtools Bioconductor package [100], the resulting SAM files were converted to BAM format and nucleotide counts performed with the pileup function for all genomic positions. The pileups were then screened for single-nucleotide polymorphisms (SNPs), requiring SNPs to exhibit a read depth between 70x and 300x for both pools, and a minor allele frequency of at least 0.2 across the pools [101]. At each of the 1,881,774 SNPs thus discovered, we then quantified the magnitude of genetic differentiation between the phenotype groups by the absolute allele frequency difference AFD [102]. SNP-specific differentiation was visualized along all chromosomes. For chromosome VI, we additionally subjected the raw AFD values to sliding window analysis to approximate the position of highest differentiation, considering windows of 10 - 100 kb, overlapping by half their width. The minimum number of SNPs required per window varied from 20 to 200, according to window width, and the median was used as window summary statistic. Since the AFD peak positions identified with the different window varied by only a few dozen kilobases, we chose 100 kb windows for final presentation. For fine-scale smoothing across the *hoxdb* region, we used locally weighted scatterplot smoothing (LOWESS) with a span of 0.0004 (relative to the whole chromosome), and a first-order polynomial.

While these analyses revealed exceptionally strong differentiation between the phenotype groups on chromosome VI, multiple segments on the pseudo-chromosome uniting all unanchored stickleback scaffolds (ChrUn) also proved highly differentiated (S2 Figure). To resolve the chromosomal origin these high-differentiation regions, we identified all chrUn contigs harboring at least one SNP with an AFD value greater than 0.4, and mapping the full contig sequence against the reference genome of the closely related ninespine stickleback *Pungitus pungitus* [103] using the BLAST services from NCBI [104]. All of these sequences consistently mapped to the ninespine stickleback chromosome homologous to the threespine stickleback chromosome VI. We thus found no evidence of any additional locus for spine reduction beside the one on chromosome VI.

### Differentiation around the hoxdb cluster in SCAD versus standard freshwater stickleback

To assess the prediction that spine-reduced SCAD stickleback exhibit strong allele frequency differentiation from completely spined standard freshwater stickleback, we sampled fin clips of 130 spine-reduced fish captured from the natural SCAD population. These tissue samples were allocated to pools of 10 individuals and subjected to DNA extraction. The 13 resulting DNA pools were then combined in equimolar proportions and sequenced as a single library in paired-end 150 bp mode to 166x median read depth on an S4 flow cell of an Illumina NovaSeq 6000 instrument.

The group for comparison comprised 100 individuals drawn from five North Uist lakes (20 individuals per lake), corresponding to the five “basic” freshwater populations described in Haenel et al. [22]. Pooled sequencing data for this reference group (paired-end 150 bp; median depth 66x) were reused from that study; full methodological details are provided therein. Following SNP ascertainment within a read-depth range of 100x-240x for the natural SCAD population and 35x-110x for the standard freshwater stickleback, AFD between the pools was calculated as for the bulk segregant analysis. Because the SCAD population is differentiated from standard freshwater stickleback at numerous loci across the genome for reasons unrelated to dorsal spine evolution [21,22], differentiation was explored around the *hoxdb* cluster only.

### RNA sequencing and gene expression analysis

To examine differences in gene expression between developing larvae from the SCAD versus marine ecotype, we sampled 30 larvae per ecotype from the second laboratory cohort at 10, 13, 16, and 19 dph (240 total individuals). This time window covered the onset of divergence between the ecotypes in cartilage and bone formation in the dorsal region, as revealed by the developmental series. The larvae were euthanized, soaked overnight at 4°C in RNAlater, and stored at -20°C until dissection. All tissue extractions were performed on the same day under a stereomicroscope, targeting the dorsal region harboring the putative second spine location using sterile scalpels and forceps treated with 70% EtOH and RNase-away (Thermo Fisher Scientific) to prevent RNA degradation. Care was taken to extract tissue samples of consistent size across individuals.

For RNA extraction, tissue collected from five individuals was combined into tissue pools, thus obtaining six biological replicate samples per developmental time point and ecotype combination. Total RNA was extracted using the PicoPure RNA Isolation Kit (Thermo Fisher Scientific), a column-based extraction method optimized for small tissue samples. Library preparation and sequencing followed standard protocols for bulk RNA sequencing at the Genomics Facility Basel (D-BSSE, ETH Zürich), including mRNA enrichment using poly(A) selection, cDNA synthesis, Illumina-compatible library construction and sequencing in 100 bp paired-end mode on two Illumina NovaSeq 6000 S4 flow cells. Sequencing depth varied across samples, ranging from 2.5 to 23.7 million reads per library (mean 11.3M, median 12.1M). Raw sequence reads were trimmed for low-quality sequences and adapters using Trimmomatic (0.39-Java-1.8; [105]), and the resulting paired-end reads were subsequently aligned to the reference genome [99] with STAR (v.2.7.3a-foss-2018b; [106]). The resulting BAM files were converted to SAM using SAMtools (v.1.20, [107]), and into a count matrix using HTSeq (v2.0.9, [108]).

Gene expression analysis was conducted using the R package DESeq2 (v.1.44.0; [109]) independently for each of the four age classes (10, 13, 16, and 19 dph). For each class, genes with very low expression levels (defined as fewer than 10 counts in fewer than three samples) were removed prior to analysis. This filtering step retained 19,037 genes at 10 dph, 18,826 at 13 dph, 18,655 at 16 dph, and 18,821 at 19 dph. Count data were normalized to counts per million reads (CPM) using the DESeq() function, and differential expression analysis was performed on the normalized counts for each age class separately. Differentially expressed genes (DEGs) between marine and SCAD stickleback were identified based on an absolute log_2_ fold change ≥ 1 and a -log_10_(*p*-value) ≥ 2. This resulted in 745 DEGs at 10 dph, 510 at 13 dph, 491 at 16 dph, and 260 at 19 dph. Because we found no substantial variation in gene expression differences between the ecotypes among the age classes, data were pooled across ages within each ecotype, thus identifying 623 DEGs. Differential expression of genes on chromosome VI was visualized for all age classes separately, flagging all DEGs located within 1 Mb to either side of the peak differentiation position between the phenotype groups in the sliding window analysis performed on the bulk segregant data.

The gene expression data were further subjected to variance stabilizing transformation to mitigate the influence of highly expressed genes. Based on the expression profiles of the top 5000 most variable genes across samples, we then performed ordination of all samples using Principal Component Analysis (PCA), thus visualizing overall differences in gene expression between the ecotypes and among age classes. In addition, we performed hierarchical clustering based on expression levels of all genome-wide DEGs based on absolute log_2_ fold change in expression (threshold: ≥ 1) and -log_10_(*p*-value) (≥ 2) using the *pheatmap* R package.

Finally, we conducted a Gene Ontology (GO) term enrichment analysis on the genome-wide DEGs (age classes pooled) to gain insights into the biological processes, cellular components, and molecular functions potentially differing in the focal tissue between the developing ecotypes. We first identified the high-confidence one-to-one zebrafish (*Danio rerio)* orthologs of all genes from the threespine stickleback genome using BioMart (Ensembl; homology type = “ortholog_one2one”, orthology confidence = 1). Subsequently, GO term enrichment analysis was performed using the orthologs of the DEGs as focal gene list, and the orthologs of all genes expressed in dorsal tissue as universe.

## Supporting information

S1 Analysis

## Acknowledgment

Field work was aided by Jamie Carmichael. Attila Ruegg, Adrian Indermaur, Deniz Ozhan, Olivia Baebler, Anja Haefeli, Laura Fritschi and Chantal Oliver helped with fish maintenance. Noëmi Behner, Nora Bisang, Philip Berner and Tino Grandchamp contributed to phenotypic analyses. Nicolas Boileau, Ana Di Pietro-Torres, Simona Doneva, Sabrina Fischer, Charlotte Huyghe, Hanna Liersch, Daniel Luescher, Maxime Policarpo and Fabrizia Ronco provided laboratory, computational and technical assistance. Sequencing was done at the Genomics Facility Basel, D-BSSE, ETH Zuerich, and calculations were performed at sciCORE, the scientific computing core facility at University of Basel (http://scicore.unibas.ch/). Novocraft shared their sequence aligner. Feedback on an early manuscript draft was provided by Katie Peichel. Financial support was provided by the Swiss National Science Foundation (SNF grant 310030_200374 to DB; SNF SINERGIA grant CRSII5_189970 to PT), the Freiwillige Akademische Gesellschaft (FAG; fellowship to CMHC) and the United Kingdom Natural Environment Research Council (NERC grant NE/R00935X/1) to ADCM. All these people and institutions are gratefully acknowledged.

## Author contributions

Conceptualization and design: DB, PT

Field work: AM, LD

Fish husbandry: CMHC

Phenotypic analyses: CMHC, TB, DB

Molecular laboratory work: CMHC, TB, AF

Bioinformatics: CMHC, DB, TB, AF

Interpretation of results: CMHC, DB, PT, TB, AF

Writing, first draft: CMHC, DB

Writing, final paper: DB, CMHC, PT

Funding: DB, PT, CMHC

## Notes

### Competing Interest Statement

The authors have declared no competing interest.

### Summary of Updates

Further phenotypic analyses have been included, such as those pertaining to the relative positioning of the spines. All figures have been updated. Candidate genes have been refined, resulting in hoxdb cluster as the main candidate responsible for the phenotypic differences.

