## Supplementary material for "Evolution of threespine stickleback dorsal spines via *hoxdb* gene regulation": S1 Analysis

**Analysis S1**

Exploration of the frequency of the derived allele in the SCAD population, and of its phenotypic effect, based on two complementary approaches using phenotype information from stickleback samples. Both approaches assumed full dominance of the derived allele causing the loss of the second dorsal spine.

The first approach combined individual-based simulations mimicking our laboratory breeding scheme over two generations with Approximate Bayesian Computation (ABC). ABC estimation targeted the observed proportion of individuals with the ancestral dorsal spine phenotype (i.e., completely spined) in the F1 and F2 hybrid (i.e., between-ecotype) cross cohorts (Fig. 1C), and estimated a frequency of the derived allele in SCAD of around 0.63 (grand median estimate with a neural network tolerance of 0.05; mode 0.62) (Figure SA). We take this estimate with some caution, given potentially imbalanced contributions of the F1 hybrid families and individuals to the F2 hybrid cohort, and given that only spine-reduced SCAD individuals were considered when initiating the laboratory lines. Based on the estimated derived allele frequency of 0.63, around 14% of the natural SCAD population are expected to exhibit the ancestral phenotype. This proportion is in qualitative agreement with the 5-10% estimated from field work in the SCAD population.

The approximately 5-10% of SCAD individuals exhibiting the ancestral completely spined phenotype formed the basis of the second approach. We here assumed that completely spined individuals in the natural SCAD population are homozygous for the recessive ancestral allele, and that the genotypes within that population are in Hardy-Weinberg equilibrium. Based on these assumptions, we estimated the derived allele to segregate in the natural SCAD population at a frequency of 0.68-0.78.

The close match in allele frequency estimates from phenotype proportions observed in independent populations (laboratory crosses vs. field samples) and different estimation strategies provides strong support for the single assumption shared between the approaches – the presence of a large-effect locus with dominance of the derived allele segregating in the SCAD population. These analyses also indicate that a substantial proportion (c. 40-50%) of the individuals in the SCAD population must be heterozygous for the ancestral and derived dorsal spine number alleles.

Further and completely independent evidence of dorsal spine reduction being caused by a dominant derived allele comes from threespine stickleback from Boulton Lake, British Columbia, Canada. In Boulton Lake, around 80% of the fish display dorsal spine reduction, while the remaining 20% reflect the typical phenotype with two major dorsal spines (Moodie & Reimchen 1973; Reimchen 1980). In the developmental genetic investigation of spine number variation in Boulton Lake by Wucherpfennig et al. (2022), a single spine-reduced Boulton Lake female was crossed with a completely spined marine male (Bodega Bay, California, USA). Although the proportions of dorsal spine number phenotypes in the resulting F1 hybrid cohort are not reported, intercrossing two completely spined F1 hybrid individuals produced only 1% (six out of 590) spine-reduced individuals among the F2 hybrid offspring. This outcome indicates that the major allele causing spine-reduction in the Boulton Lake population is dominant; the completely spined F1 hybrid individuals used for generating the F2 hybrid cohort must have been homozygous for the recessive ancestral allele associated with the complete dorsal spine phenotype. The crossing design of the Wucherpfenning et al. (2022) investigation thus appears to have excluded the major allele underlying the naturally occurring dorsal spine reduction in Boulton Lake, and instead focused on an alternative allele *increasing* dorsal spine number beyond the typical completely spined phenotype (3.4% of their F2 hybrid cohort displayed three major dorsal spines instead of two).

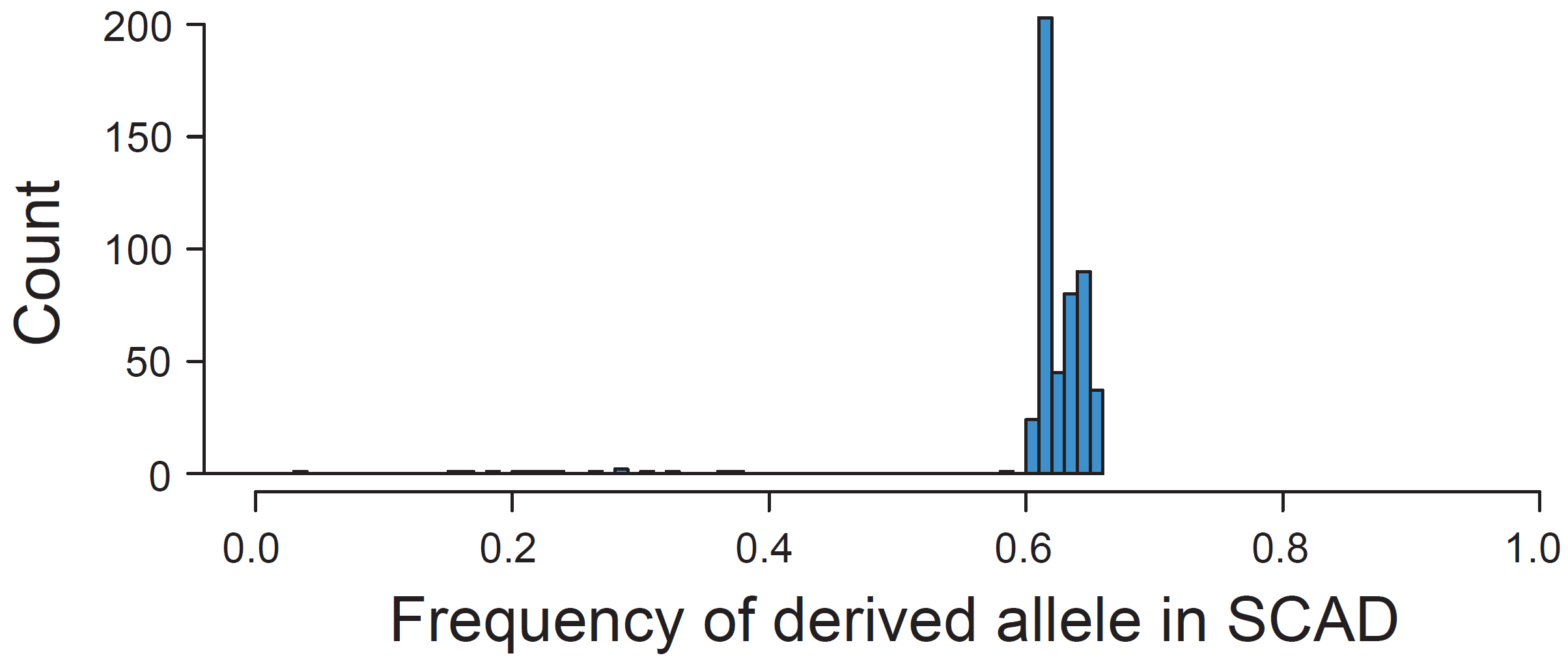

**Figure SA.** **Frequency and phenotypic effect of the spine reduction allele.** Distribution of median estimates of the frequency of the dominant derived allele in the SCAD population across 500 ABC estimation runs. The grand median is 0.63 (mode 0.62).
