## Supplementary material for "Evolution of threespine stickleback dorsal spines via *hoxdb* gene regulation": S1 Figure

**Figure S1**

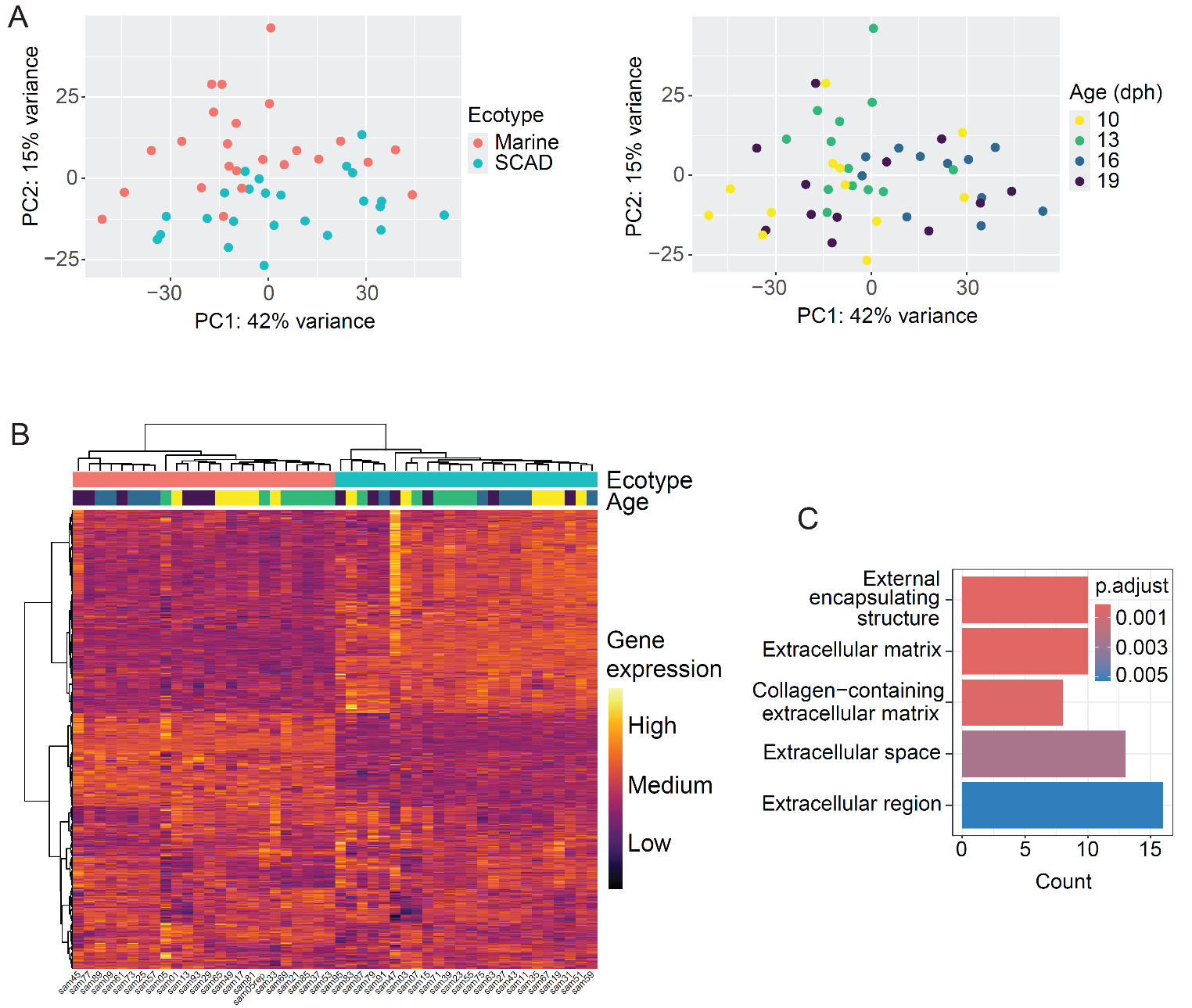

**Figure S1. Global gene expression.** **A)** Position of individual samples, each representing a tissue pool of five fish, along the first two principal components (PC1, PC2) extracted from gene expression variation in the top 5000 variable genes. The samples are color-coded by source population (ecotype; left) and age (days post hatching; right). **B)** Heatmap of gene expression Z-scores, with hierarchical clustering, computed for the 623 differentially expressed genes (DEGs; rows) between the marine and SCAD samples (data pooled across the four age classes within each ecotype). The labels below the graph indicate the sample names. **C)** Gene Ontology terms associated with Cellular Component (CC) enriched among the DEGs between marine and SCAD stickleback.
