## Supplementary material for "Evolution of threespine stickleback dorsal spines via *hoxdb* gene regulation": S2 Figure

**Figure S2**

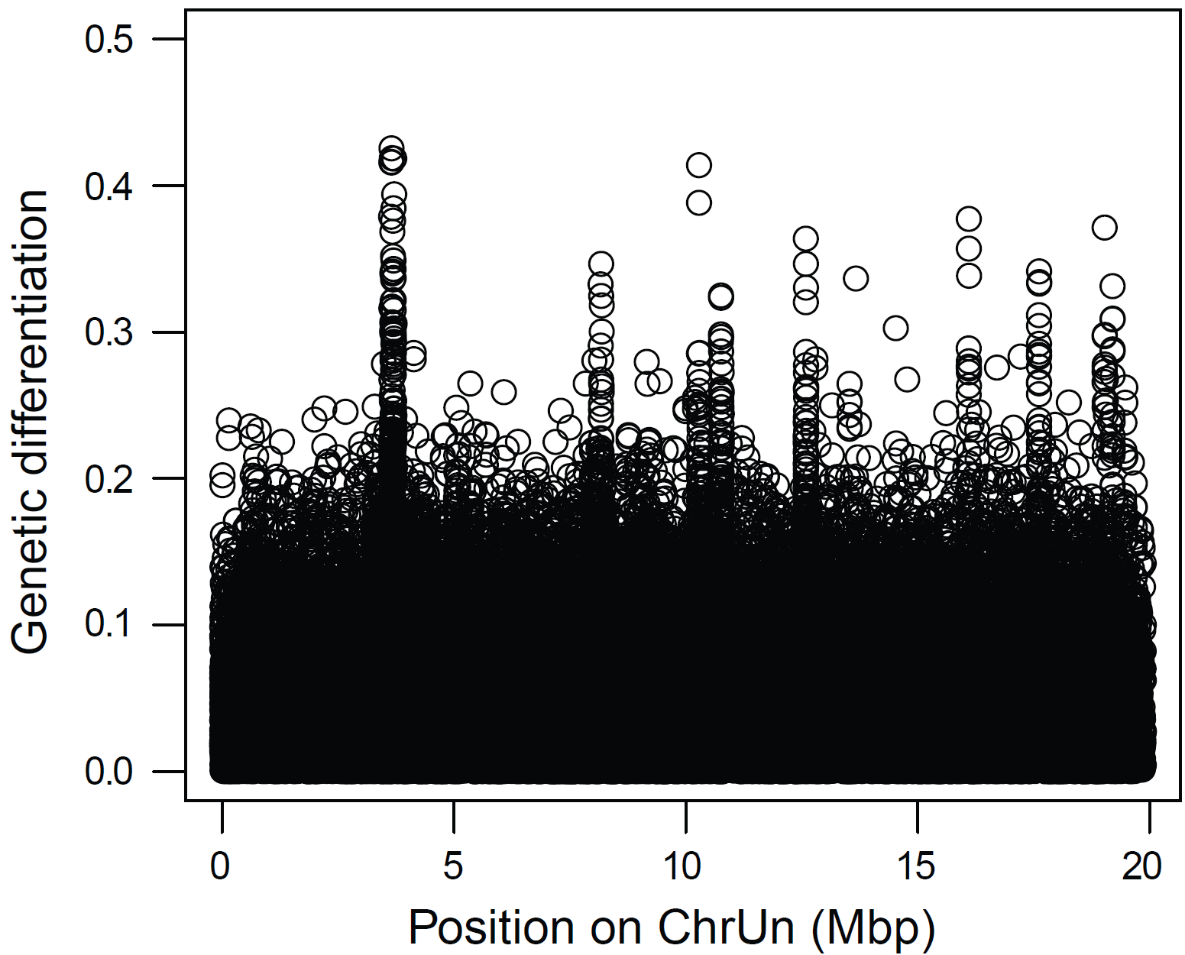

**Figure S2. Chromosomal origin of high-differentiation loci on ChrUn.** Close-up of the pseudo-chromosome ChrUn harboring contigs with high genetic differentiation (AFD) between the marine and SCAD population in the bulk segregant experiment. All contigs with strong differentiation proved to belong to chromosome VI.
