## Supplementary material for "Evolution of threespine stickleback dorsal spines via *hoxdb* gene regulation": S1 Methods

**Methods S1.** **Bone staining protocol used for the bulk segregant analysis.**

**Alizarin Red bone staining protocol**

1. Place the specimen in Formalin 10% for 24 hours.

2. Transfer each individual to Alizarin Red and let submerged for 24 hours.

3. Rinse the specimen with distilled pure water to remove excess of Alizarin Red.

4. Transfer each individual to KOH solution and shake it vigorously. Leave it soaking for 4 hours.

5. Substitute the KOH solution for distilled pure water and let it soak overnight.

**Required solutions**

Formalin 10%

1. Pour 800 ml of distilled pure water in a glass beaker with magnetic stirrer.

2. Add 40g of Paraformaldehyde and 8g of Phosphate Buffered Saline.

3. Add roughly 28 drops of NaOH 1M with Pasteur pipette and wait.

4. Measure the translucent solution with calibrated pH meter.

5. Adjust the pH with the fuming HCl as follows: pour a small drop into your solution and let it mix well for some seconds and then re-measure pH.

6. Repeat step 5 Until pH is around 7.

7. Add 200 ml of distilled pure water and store.

Alizarin Red

1. Pour 1800 ml distilled pure water in a glass beaker with magnetic stirrer.

2. Add 4.2 g KOH chips.

3. Add 0.4 g Alizarin Red powder. The solution turns dark purple. Allow it to mix for 2 minutes.

4. Add 200 ml distilled pure water.

KOH solution

1. Pour 1800 ml distilled pure water in a glass beaker with magnetic stirrer.

2. Add 0.2 g KOH chips. Wait until completely dissolved.

3. Add 200 ml distilled pure water.
